# Sisyphus at the splenic flexure: sidedness in colorectal cancer

**DOI:** 10.64898/2026.11.07.761937

**Authors:** Liangze Ke, Michael Cecchini, Vincent L. Cannataro, Jeffrey P. Townsend

**Affiliations:** Department of Biostatistics, Yale School of Public Health, New Haven, CT, USA; Program in Developmental Therapeutics, Yale Cancer Center, New Haven, CT, USA; Department of Medicine, Yale School of Medicine, New Haven, CT, USA; Department of Biology, Emmanuel College, Boston, MA, USA; Program in Genetics, Genomics, and Epigenetics, Yale Cancer Center, New Haven, CT, USA; Department of Ecology and Evolutionary Biology, Yale University, New Haven, CT, USA

**Keywords:** colorectal cancer, colorectal sidedness, driver genes, somatic evolution, mutational signatures

## Abstract

Colorectal cancer is divided into left and right to guide prognosis and therapy, yet a precise definition of left vs right sided colorectal cancer is still not fully agreed upon. The published conventions place it anywhere between the cecum and the rectosigmoid junction. The dichotomy assumes side homogeneity, i.e., that colorectal subsites within a side add limited information once the side is known. That assumption has been examined primarily through mutation prevalence, a quantity that confounds how often a driver arises with how strongly it is selected once it has. Analyzing 1,027 whole-exome and whole-genome sequenced colorectal tumors, we decomposed the prevalence of six driver groups into mutation supply and cancer effect, and asked whether a binary side label captures how each of their prevalence, mutation supply, and cancer effect vary along the colorectum. Mostly it does: the mutational input behaves as two blocks, and even the more than thirty–fold strengthening of cancer effect on *BRAF* V600E from sigmoid colon to cecum is adequately captured by side. But *APC* shows nominal evidence against side homogeneity in both prevalence and cancer effect while showing little net left–right difference, and *PIK3CA* H1047 rejects it in supply. The evidence for finer resolution is real but partial, and the failures do not announce themselves. We recommend additional research recording primary tumor location at the finest practicable resolution—at minimum, International Classification of Diseases for Oncology, 3rd Edition (ICD-O-3) topography. Such well-annotated datasets would enable the left–right boundary to be a reversible analytic choice rather than an irreversible annotation.

## Main Text

Primary-tumor location is among the most robust and clinically actionable variables in colorectal cancer (CRC). Its conventional representation by sidedness—left or right—captures numerous molecular and clinical contrasts and carries substantial prognostic weight (**Table 1**). However, clinical utility does not establish sidedness as a natural biological unit. Collapsing the colorectum into two bins is sufficient only if finer positioning within each side provides no promise of etiological, molecular, prognostic, or predictive information. Where the left–right boundary is drawn is a matter of convention rather than consensus. Current guidelines draw it at the splenic flexure, classifying tumors from the flexure through the rectum as left-sided (Figure 1) [1]; they use sidedness to guide first-line therapy yet describe primary-tumor location as only “surrogate information” for underlying biology [1]. They use sidedness to guide first-line therapy, yet describe primary-tumor location as “surrogate information” for underlying biology [1]. The usual biological rationale for sidedness is embryological: the colorectum develops from midgut and hindgut segments, each supplied by a distinct aortic branch, with their boundary where the two blood supplies meet [2]. However, that embryological transition arises two-thirds of the way along the transverse colon—roughly 15 cm proximal to the typically used clinical boundary (Figure 1), [3, 4]. Some investigators use the embryological border instead[5, 6]. When two biologically plausible conventions can assign the same tumor to opposite sides, sidedness is not an intrinsic property of the tumor but a categorical label imposed on continuous anatomy.

**Fig. 1.**
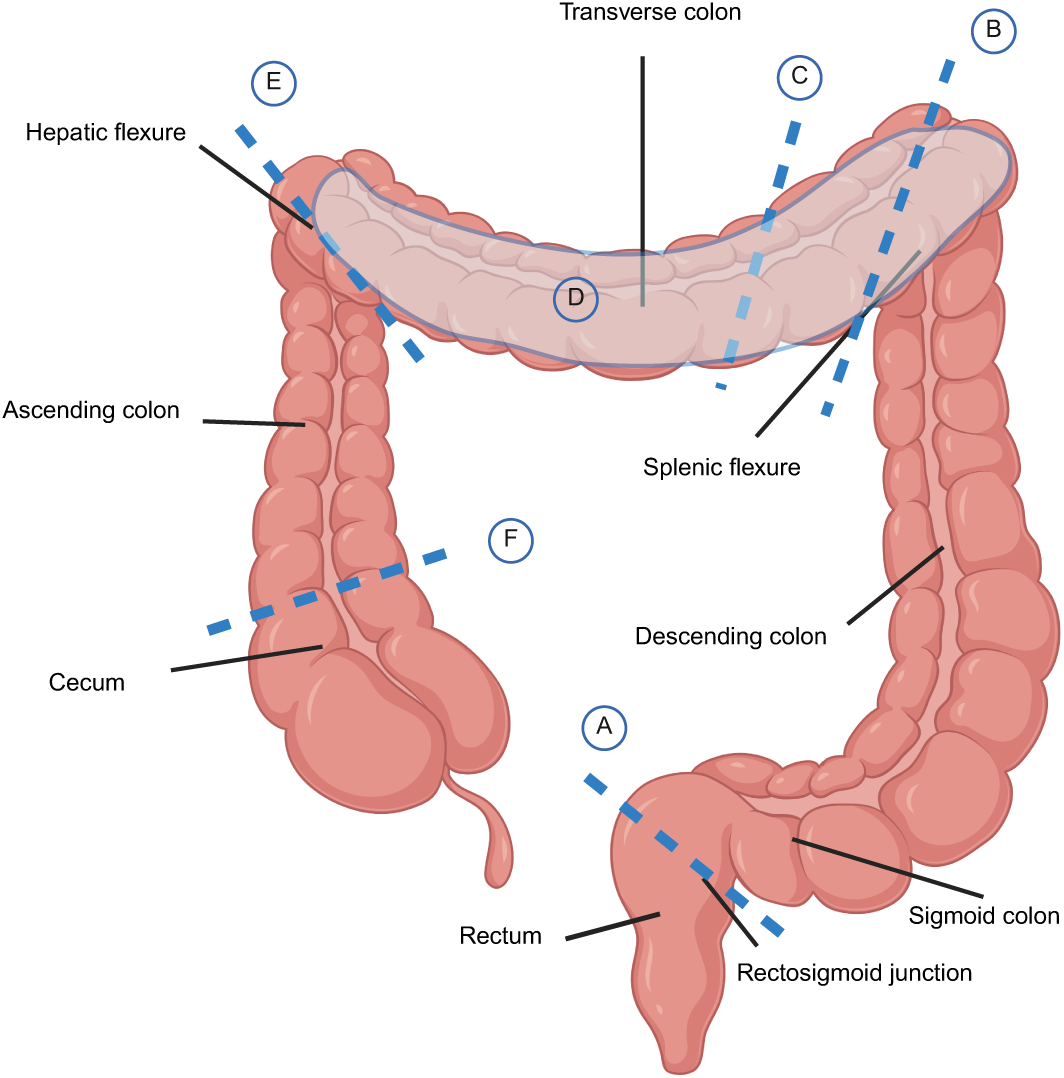
Competing anatomical definitions of colorectal sidedness. Schematic of the large bowel showing anatomical boundaries and contested regions (dashed lines, **A**–**F**) that have been used or proposed, singly or in combination, to classify primary tumors as “left-” or “right-sided.” Proceeding from the rectum (”leftmost” anatomy) to cecum (”rightmost” anatomy): **A** the sigmoid– rectal junction, used to distinguish rectal from the colonic tumors [10, 15, 16]; **B** the proximal margin of the splenic flexure, corresponding to the current clinical convention [1], which assigns splenic-flexure tumors to the left, while other definitions assign them to the right or exclude them from both sides[3, 10–13]; **C** the embryologic junction between midgut- and hindgut-derived bowel within the transverse colon, marks the developmental boundary commonly invoked to justify sidedness[5, 6]; **D** the transverse colon and both flexures, constituting a mid-colonic “gray area” whose classification varies among schemes[13]; **E** the hepatic-flexure boundary, which has been paired with the splenic flexure boundary to exclude transverse-colon tumors from both sides [60]; and **F** the cecum-ascending colon junction, which has been used to distinguish cecal tumors from the remainder of the “right” colon [17]. These many boundary definitions mean that a single primary tumor location can be classified as left-sided, right-sided, or neither, depending on the definition applied.

**Table 1.** Clinical and molecular contrasts between left- and right-sided colorectal cancer.

| Feature | Left-sided | Right-sided |
|---|---|---|
| Embryonic origin | Hindgut (inferior mesenteric artery) | Midgut (superior mesenteric artery) |
| Prognosis<sup>a</sup> | Better; median OS 33.3 months | Worse; median OS 19.4 months |
| Anti-EGFR benefit<sup>b</sup> | Most of the benefit | Little to none |
| Genomic instability<sup>c</sup> | Chromosomal instability | Microsatellite instability (MSI-high/dMMR) |
| *BRAF* and methylation<sup>c</sup> | *BRAF* wild-type, CIMP-low | *BRAF*-mutant, CIMP-high |
| Precursor pathway<sup>c</sup> | Conventional adenoma–carcinoma sequence | Serrated pathway (sessile serrated lesions) |

<sup>a</sup>Median overall survival in metastatic CRC from CALGB/SWOG 80405 [8]; the right-sided disadvantage persists across stages [10, 61]. <sup>b</sup>In *RAS* wild-type metastatic CRC [1, 7, 9, 25]. <sup>c</sup>Features are enriched in, not exclusive to, the indicated side [24, 31, 32, 62]. Abbreviations: CIMP, CpG island methylator phenotype; CRC, colorectal cancer; dMMR, mismatch repair deficient; EGFR, epidermal growth factor receptor; MSI, microsatellite instability; OS, overall survival.

This inconsistency is embedded in the clinical evidence. CRYSTAL and FIRE-3 trials divided at the splenic flexure and classified the transverse colon as right-sided [7], whereas CALGB/SWOG 80405 used both flexures as boundaries and placed the transverse colon on neither side [8]. A six-trial pooled analysis reconciled these differences only by excluding transverse-colon tumors from one trial while treating them as right-sided in the other five [9]. The splenic flexure is assigned to the left by most schemes, to the right by others [3, 10]. Still others exclude it from both sides [11–14]—implicitly dissecting the colorectum into three. The poles are similarly unstable: the rectum has been separated from the left colon because of copy-number, mutational, and transcriptional distinctions [10, 15, 16], and the cecum from the right colon because its prognosis differs from that of the ascending colon[17]. Applying published definitions to a single cohort, Lansom et al. [13] identified seven divisions, with boundaries ranging from the ascending colon to the rectosigmoid junction. Nor is the contested region negligible: the transverse colon and both flexures constitute the longest and most variable portion of the large bowel, with a mean length of 47 cm [18]. Across this distance, the same tumor can be assigned to different sides, and different biological, prognostic, and therapeutic interpretations solely through the choice of scheme. Sidedness thus converts continuous anatomy into a scheme-dependent variable, which may not capture the full biology of the disease.

For more than two decades, findings challenging within-side homogeneity have usually prompted investigators to relocate the boundary or exclude a discordant segment rather than test whether any binary partition is adequate. Proposed divisions derive variously from embryological and vascular supply [2, 5, 6, 19]; survival analyses selecting the cutpoint that maximizes prognostic separation [17, 20–23]; molecular and histopathological profiles [13, 15, 16, 24]; treatment-response analyses that exclude nonconforming segments [7–9, 25]; and expert anatomical consensus [26, 27]. These resulting cutpoints are each internally coherent. However, they are mutually incompatible. Each can restore local homogeneity for one subsite, endpoint, or cohort while leaving untested the premise of two homogeneous sides. Each new discordance can prompt another boundary revision. A different site or endpoint can then expose another discordance and prompt another cutpoint, perpetuating a Sisyphean cycle of boundary revision.

Yamauchi et al. were among the first to step out of this Sisyphean cycle, examining molecular variation across colorectal subsites without imposing a boundary in advance, and arguing that the colon is a continuum [28, 29]. Loree et al. subsequently demonstrated substantial heterogeneity within both conventional sides: among right-sided tumors, RAS mutation prevalence declined from 70% in the cecum to 43% at the hepatic flexure while *BRAF* V600E prevalence increased from 10% to 22% [20]. Moreover, Cornish et al. extended this spatial view through whole-genome sequencing of colorectal cancers across nine colorectal subsites, showing that microsatellite-stable primary tumors exhibited coordinated gradients in variant burden, mutational signatures, driver frequencies, and genomic subgroups. Moving distally, SNV and indel burdens declined, structural-variant and copy-number burdens increased, colibactin-associated mutagenesis became more prominent, *TP53* mutation prevalence increased, and *KRAS*, *PIK3CA*, and *AMER1* mutation prevalence decreased [30]. Together, these studies suggest an escape from the Sisyphean cycle: not another boundary, but a positional map capable of resolving how mutational processes, driver prevalence, and clonal selection vary along the colorectal continuum.

Whether or not adhering to this two-sides dichotomy, authors generally examined prevalence of expression-defined consensus molecular subtypes [31] and of mutations in *TP53*, *KRAS*, *BRAF* V600E, and other genes [20, 28, 30, 30, 32]. Prevalence, however, is the compound outcome of two somatic evolutionary forces: mutation supply and clonal selection. Under the strong-selection–weak-mutation model used to estimate cancer effect, the expected flux of successful driver lineages equals the neutral mutation rate multiplied by the cancer effect, or scaled selection coefficient [33]. For each gene in each tumor, the neutral mutation rate—the rate expected without selection—is calculated from signature-weighted trinucleotide rates adjusted for local gene mutability. Cancer effect measures how strongly selection for cellular survival and proliferation amplifies that expectation: a value of one denotes neutrality, whereas values greater than one indicate positive selection [34]. A driver may be enriched in the right colon because it arises more readily there, as when proximally concentrated microsatellite instability raises supply, because it confers a greater selective advantage there, or both; prevalence alone cannot distinguish these mechanisms. Mutation supply most directly informs prevention, whereas selection most directly informs treatment.

Here we question whether sidedness exhausts the spatial information in driver prevalence, neutral mutation supply, and cancer effect. Primary colorectal adeno-carcinoma whole-exome or -genome somatic mutation calls that were annotated with International Classification of Diseases for Oncology, Third Edition (ICD-O-3) anatomical subsites [35] [36] were obtained from the Catalogue of Somatic Mutations in Cancer (COSMIC) v104. The primary analytic cohort comprised 1,027 tumors localized to the rectum (C20; *n* = 382; 37.2%), rectosigmoid (C19; 75; 7.3%), sigmoid colon (C18.7; 159; 15.5%), descending colon (C18.6; 19; 1.9%), splenic flexure (C18.5; 8; 0.8%), transverse colon (C18.4; 32; 3.1%), hepatic flexure (C18.3; 14; 1.4%), ascending colon (C18.2; 130; 12.7%), and cecum (C18.0; 208; 20.3%) (**Supplementary Information**). Under the clinically used dichotomy [1], 643 tumors were left-sided and 384 were right-sided.

Cohort assembly exposed a data limitation central to the sidedness problem. Many source studies reported only investigator-defined side categories rather than ICD-O-3 colorectal subsites [35]. Once subsite is collapsed to side, the original position cannot be recovered. The same problem is also reported in a 2,023-genome national cohort [30]. This irreversible aggregation, together with genuinely low mid-bowel incidence (roughly 10% of CRC[37]), likely contributed to uneven representation of tumors along the colorectal axis in our dataset. The segments most affected by competing boundary definitions—the descending colon, splenic flexure, transverse colon, and hepatic flexure—contributed only 73 assignable tumors (7.1%); others had been merged with a neighboring side under unspecified criteria or labeled simply as “other”. Thus, the annotation practices that operationalize sidedness have also limited the data available to evaluate it.

For each tumor, single-base substitution spectra were deconvolved into COSMIC v3.4 mutational signatures [38, 39] and consolidated into nine etiologic categories [40], yielding the proportional contribution of each process at every colorectal subsite (Figure 2). Sidedness captured two robust opposed patterns. Defective mismatch repair (dMMR) accounted for 4.7% of mutations in left-sided tumors and 21.6% in right-sided tumors, a 4.6-fold enrichment; the ranges of dMMR among subsites on the left (all *≤* 5.5%) did not overlap with the ranges of dMMR among subsites on the right (*≥* 20.2%). This right-sided enrichment on the right accords with sporadic dMMR arising via *MLH1* promoter hypermethylation in CIMP-high tumors derived from sessile serrated lesions, which occur disproportionately in the right colon [41, 42]. APOBEC-associated mutagenesis exhibited the inverse mutational pattern, contributing 2.8% of mutations on the left and 1.4% on the right, again with non-overlapping subsite ranges (left, *≥* 2.4%; right *≤* 1.7%). This side-level contrast is consistent with recent observations across 2,023 colorectal cancers, in which the APOBEC-associated trinucleotide mutation signatures SBS2 and 13 were enriched in microsatellite-stable (MSS) but not in microsatellite-instable (MSI) tumors. Because MSI tumors are concentrated on the right-sided colorectum, this subtype difference predicts the left-sided APOBEC enrichment observed here. Within MSS tumors, however, neither SNS 2 nor SBS13 showed significant anatomical variation, suggesting that much of the side-level contrast may correspond to the spatial distribution of MSI rather than an independent anatomical gradient [30]. Additional research should be performed to ascertain if the opposite sidedness of APOBEC-associated mutations is driven entirely by MSI, or may additionally reflect APOBEC3 antiviral activity [43, 44], cyclic hypoxia and the ensuing replication catastrophe [45–47], synergy between fragile histidine triad loss and APOBEC3B overexpression [48], or other mechanisms [49, 50]. Other processes conformed less closely to sidedness: defective homologous recombination was more prominent on the left but also varied from subsite to subsite within sides, whereas clock-like SBS5, non-actionable, and unknown signatures varied significantly along the colorectum without a clear left-right pattern, consistent with graded proximal-to-distal declines in SBS1 and SBS5 activity among microsatellite-stable tumors [30]. Thus, sidedness captures prominent differences in mutational input but not its full spatial structure.

**Fig. 2.**
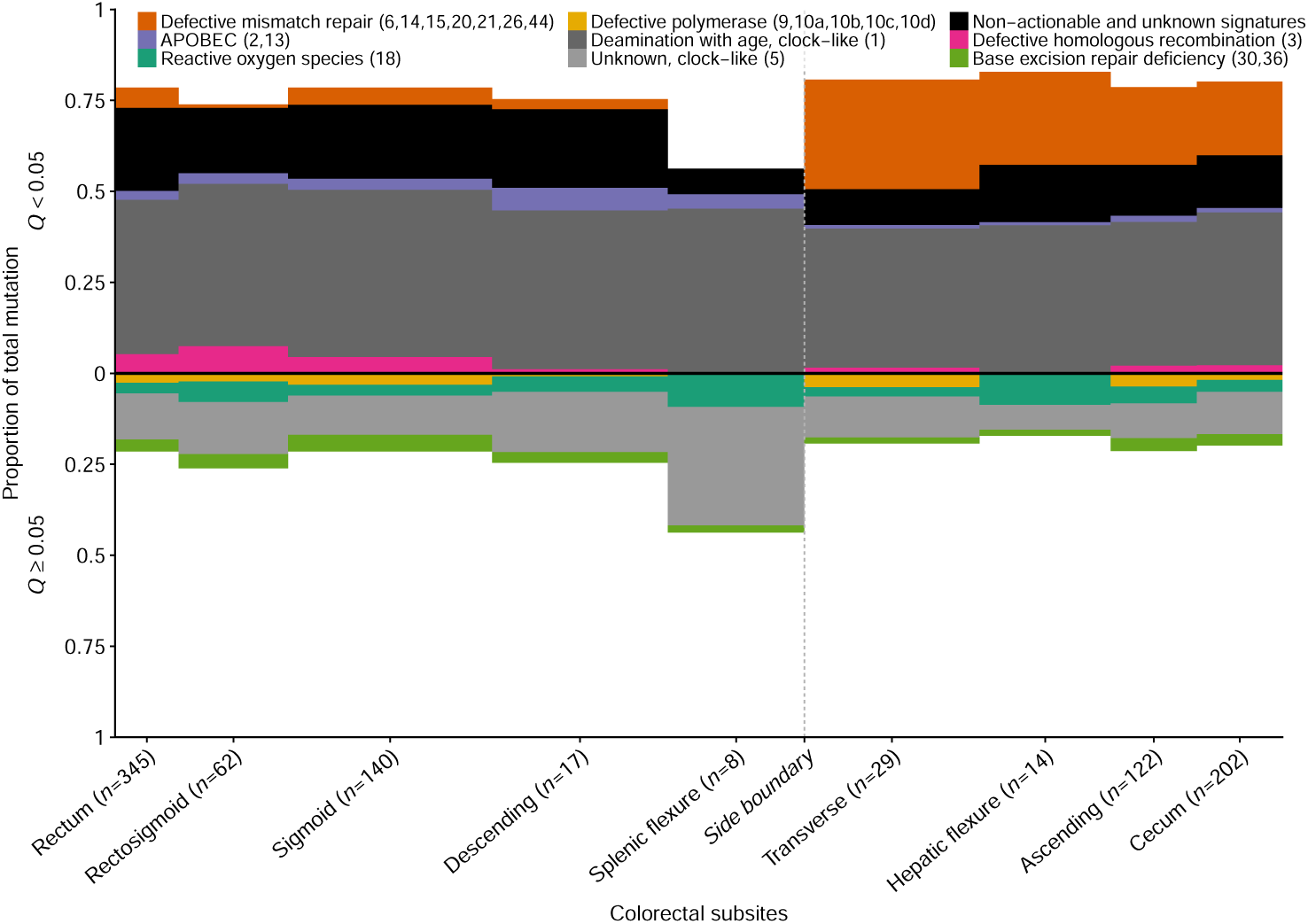
Positional variation in mutational-process contributions across colorectal subsites. Single-base substitutions from *n* = 939 whole-exome and -genome sequenced colorectal tumors with *≥* 50 single nucleotide variations were decomposed into Catalogue of Somatic Mutations in Cancer (COSMIC) version 3.4 signatures and grouped into nine etiologic processes. For each of nine anatomical subsites defined by the International Classification of Diseases for Oncology, Third Edition (ICD-O-3), each bar segment shows that etiology group’s proportion of total mutation, averaged across patients within the subsite; the absolute segment heights above and below the baseline sum to 1. Subsites are ordered from rectum to cecum, with horizontal widths proportional to their approximate anatomical lengths [18]. Etiologic groups whose contributions differed significantly among subsites are plotted above the baseline, stacked upwards in order of decreasing *Q* values (likelihood-ratio test with Benjamini–Hochberg adjustment for 9 tests, *Q <* 0.05); the positive contributions of remaining etiologies that are not significantly different across subsites are plotted below, stacked downwards in order of increasing *P* values.

Tumor-wide mutational-process contributions, however, do not equate with contribution to cancer. Selection acts on specific variants, so a process that dominates the overall mutation spectrum need not generate the mutations contributing most strongly to tumorigenesis. The two quantities can diverge sharply: dMMR-associated processes account for 21.6% of all mutagenesis in right-sided tumors, but they account for 58–91% of the cancer effect of *BRAF* V600E and only 6–13% of that of *TP53* across right-colon subsites. This divergence reflects driver-specific mutational targets, whose sites occupy distinct trinucleotide contexts; the same mixture of processes therefore generates mutations in different drivers at different relative rates. We therefore decomposed cancer effect into the fractions attributable to individual mutational processes, separately for each of the six drivers sufficiently prevalent for subsite-resolved estimation[40] (Figure 3A; Supplementary Information).

**Fig. 3.**
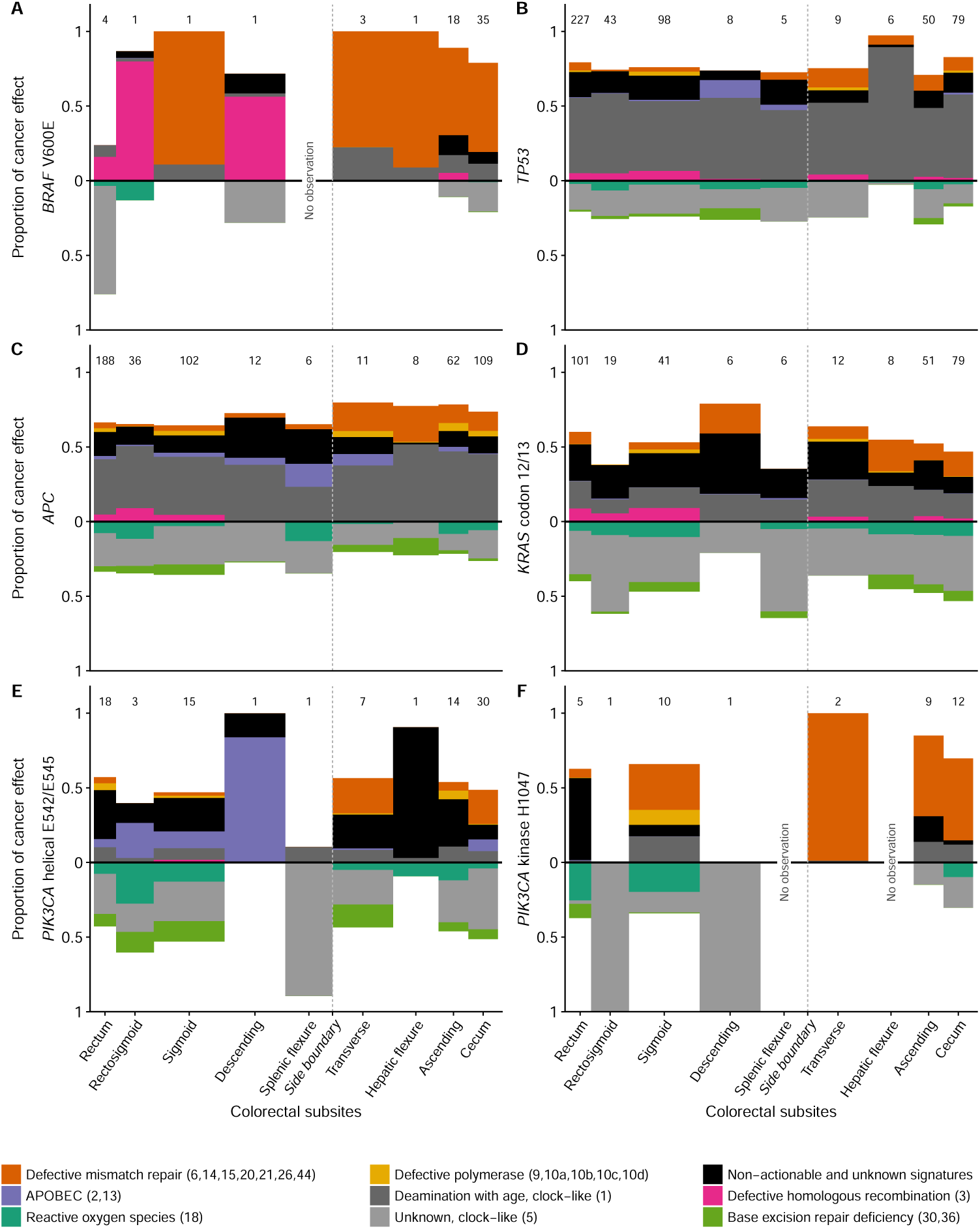
Mutational-process attribution of driver-specific cancer effects across colorectal subsites. Analyzing *n* = 1027 whole-exome and -genome sequenced tumors for the cancer effect—the scaled selection coefficient inferred from driver prevalence after accounting for tumor-specific neutral mutation supply—of **A** *BRAF* V600E, **B** *TP53*, **C** *APC*, **D** *KRAS* codons 12/13, **E** the *PIK3CA* E542/E545 helical-domain hotspots, and **F** the *PIK3CA* H1047 kinase-domain hotspot, partitioned among the mutational processes capable of generating the variants and expressed as proportional contributions summing to one. Colors represent etiologic groups derived from Catalogue of Somatic Mutations in Cancer (COSMIC) version 3.4 single-base-substitution signatures. Etiologic groups are stacked in the same order as in Figure 2. Subsites are ordered from rectum to cecum, and numbers above the bars indicate the number of tumors carrying the corresponding driver at each subsite. The dashed vertical line marks the clinical left–right boundary at the splenic flexure, with splenic-flexure tumors classified as left-sided.[1].

The clearest side-specific difference in this site-specific attribution involved the dMMR contribution to *BRAF* V600E. Among the four left-sided subsites with at least one *BRAF* V600E tumor, the dMMR-attributed fraction of cancer effect was 0% at three (with the sigmoid colon as a marked exception at 89.1%). Across all four right-sided subsites, this fraction ranged from 58.4–91.1% (Figure 3A). This pattern accords with the co-occurrence of *BRAF* V600E and *MLH1* silencing within the right-enriched serrated pathway [41, 42]. Greater right-sided dMMR attribution was also evident for *TP53*, *APC*, and *PIK3CA* H1047, and less consistently for *PIK3CA* E542/E545 (Figure 3B,C,E,F). APOBEC-associated mutagenesis showed the opposite sidedness, accounting for 11.1% of cancer effect of mutations on the left and 4.7% on the right, specifically for the *PIK3CA* helical-domain hotspots E542/E545. No other driver group showed appreciable APOBEC attribution. This specificity is consistent with sequence context: E542K and E545K are C *→* T substitutions in the APOBEC-preferred TCA context on the pyrimidine-oriented strand [38, 44]. HPV-associated APOBEC activation provides a mechanistic precedent in other cancers [51–53], its provenance deserves additional research attention. Resolving mutational-process contributions driver by driver confers additional mechanistic specificity to broad side-level etiologic patterns.

Having observed substantial variance in the processes capable of generating each driver, we quantified the degree to which mutation supply varied across colorectal subsites. Driver-specific neutral mutation rates, summed across underlying mutational processes, were strikingly stable (Figure 4)[40]: All six varied by less than twofold; for four, the full range was less than 25% of the mean: *TP53* (11.9%), *APC* (20.1%), *PIK3CA* E542/E545 (17.8%), and *PIK3CA* H1047 (11.8%). *BRAF* V600E (Figure 4A) varied the most, by 1.8-fold, declining from its maximum in the rectum to its minimum in the transverse colon before recovering modestly toward the cecum. Thus, the right-sided enrichment of *BRAF* V600E cannot be explained by greater mutation supply. Indeed, assessment of mutational processes alone would predict the opposite sidedness for *BRAF* V600E mutations.

**Fig. 4.**
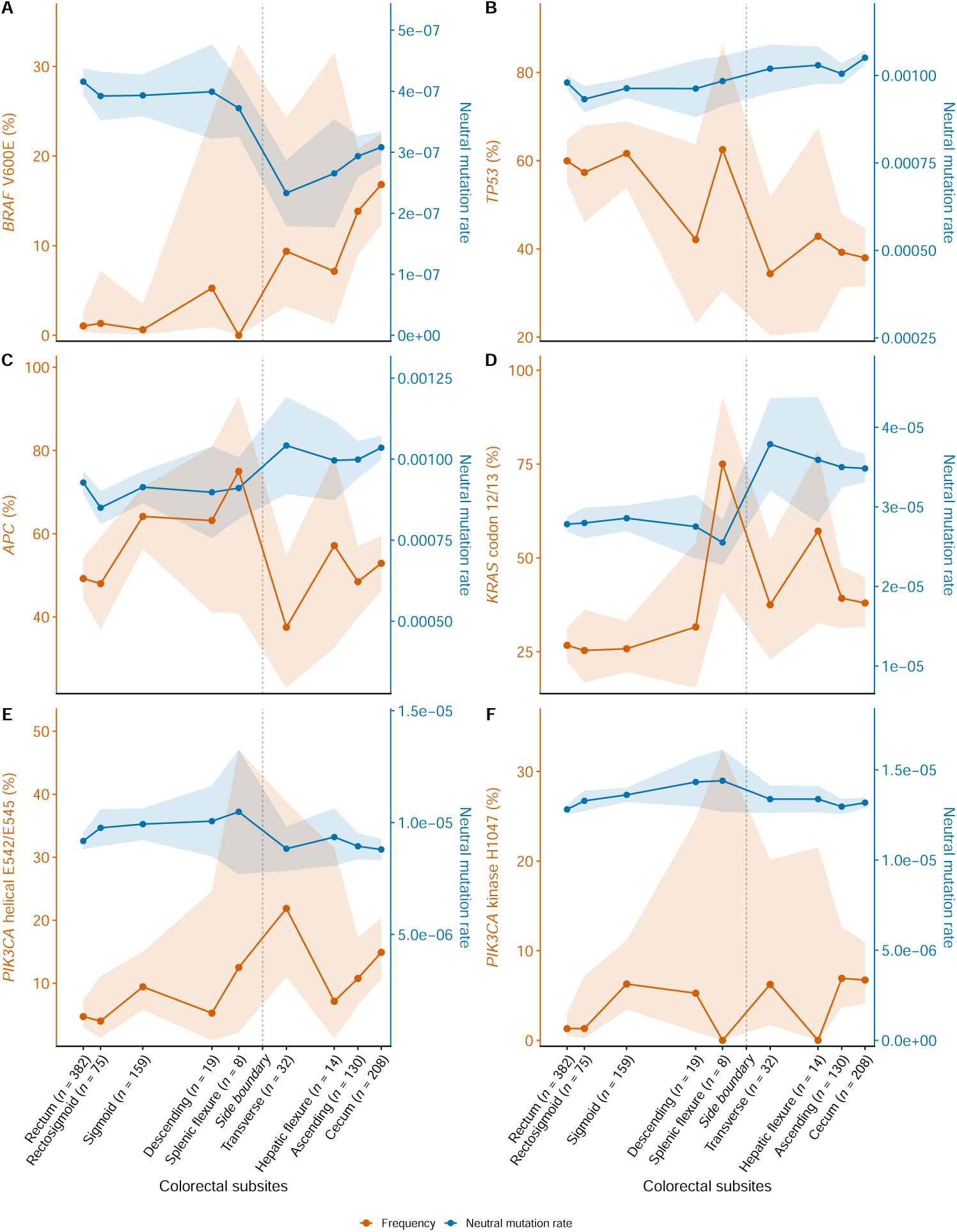
Driver prevalence and neutral mutation supply across colorectal subsites. Prevalence and tumor-specific neutral mutation supply are traced across nine primary-tumor subsites from *n* = 1027 whole-exome and -genome sequenced tumors, ordered from rectum to cecum, for **A** *BRAF* V600E, **B** *TP53*, **C** *APC*, **D** *KRAS* codons 12/13, **E**, *PIK3CA* helical E542/E545; and **F**, *PIK3CA* kinase H1047 plotted against primary tumor location. Driver mutation prevalence (orange, left axis; shaded band, Wilson 95% CI) is the proportion of tumors carrying the specified driver. Neutral mutation rate (blue, right axis; shaded band, *±*1.96 SE) is mean tumor-specific neutral mutation rate at the driver site or sites, which corresponds to the expected rate at which the driver would arise in the tumor in the absence of selection. The dashed vertical line marks the clinical left–right boundary at the splenic flexure, with splenic flexure tumors classified as left-sided[1].

The observed spatial differences in driver prevalence therefore appear to arise primarily from what happens after the mutations are first generated: location-dependent variation in clonal selection. Across subsites in which the relevant driver was observed, estimated cancer effects varied by as much as 36-fold (Figure 5A), whereas neutral mutation rates varied by no more than 1.8-fold. The clearest opposing gradients again involved *BRAF* V600E and *TP53*. The cancer effect of *BRAF* V600E in the colorectum increased by more than an order of magnitude, from 1.6*×*10^4^ in the sigmoid colon to 5.7 *×* 10^5^ in the cecum, rising within as well as between conventionally defined sides (Figure 5). In contrast, the cancer effect of *TP53* mutations declined by approximately half from the rectum to the cecum (Figure 5B)

**Fig. 5.**
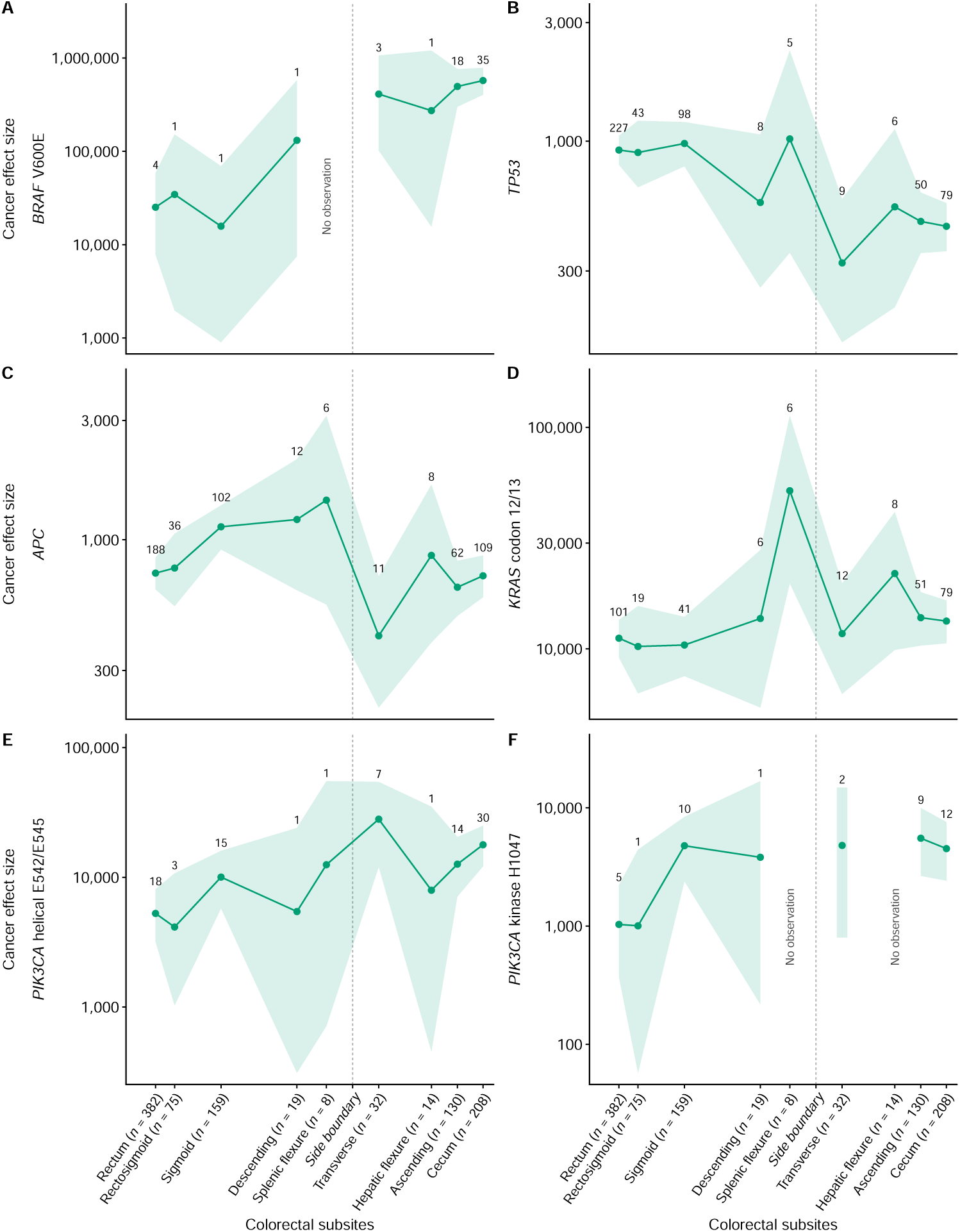
Driver-specific cancer effects across colorectal subsites. Cancer effects—the scaled selection coefficient inferred from driver prevalence accounting for neutral mutation supply—were estimated separately at each of nine primary-tumor subsites for a total of *n* = 1027 whole-exome and -genome sequenced tumors, ordered from rectum to cecum, for (**A** *BRAF* V600E, **B** *TP53*, **C** *APC*, **D** *KRAS* codon 12/13, **E** *PIK3CA* E542/E545 helical-domain hotspots, and **F** *PIK3CA* H1047 kinase-domain hotspot. Points denote subsite-specific cancer-effect estimates, connected by lines and shaded 95% confidence intervals. Numbers above points provide the count of tumors carrying the corresponding driver at each subsite. The dashed vertical line marks the clinical left–right boundary at the splenic flexure, with splenic flexure tumors classified as left-sided [1].

The markedly different magnitudes of these estimates reflect the drivers’ different mutational target sizes. The *BRAF* V600E substitution is primarily generated by a single-base substitution yielding a constitutively active, *RAS* -independent monomeric kinase [54, 55]. Its neutral mutation rate is the lowest among the six driver groups. Consequently, its frequent recurrence implies exceptionally strong clonal selection[33]. In contrast, the *TP53* driver group comprises hundreds of truncating and deleterious substitutions distributed across the gene, predominantly within its large DNA-binding domain. Their aggregate neutral mutation supply is approximately 2,700-fold greater than that of *BRAF* V600E. Therefore, a much smaller shared selection coefficient can account for the observed aggregate prevalence.

Together with the broader prevalence gradients, these driver-specific patterns suggest that cancer effects vary among primary-tumor subsites beyond what can be conveyed by a distinction between left and right. Across all six driver groups, point estimates varied substantially and sometimes nonmonotonically among subsites within each side. However, the small numbers of tumors carrying a given driver at several subsites yielded imprecise estimates with wide confidence intervals. To determine whether current data could resolve positional variation beyond sidedness, we compared side-only models, estimating one value for left-sided tumors and another for right-sided tumors, with subsite-resolved models, estimating separate values for each of nine annotated subsites. Splenic-flexure tumors were classified as left-sided in accordance with current clinical convention [1]. Because the models are nested, a likelihood-ratio test asks whether subsite explains variation beyond side; with seven additional parameters, the test is conservative to test sidedness. We first applied this assessment to driver prevalence.

The likelihood of the observed prevalence of each driver in anatomical locations

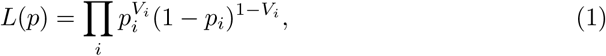

where *V_i_* = 1 if tumor *i* carries the driver and 0 otherwise, and *p_i_* is the probability of carrying the driver in that location. The side-only and subsite-resolved models were

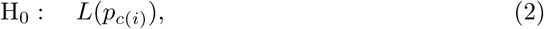

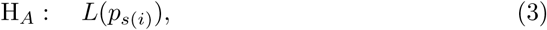

where *c*(*i*) *∈ {L, R}* identifies the side of tumor *i*, so that *p_c_*_(*i*)_ takes one value per side, and *s*(*i*) *∈* {1, . . ., 9} identifies the subsite, so that *p_s_*_(*i*)_ takes one value per subsite. Maximized likelihoods were compared using a likelihood-ratio test with seven degrees of freedom.

This conservative test on limited data did not reject the side-only model for five of six drivers (Table 2). *APC* prevalence showed nominal evidence of subsite variation beyond sidedness (*P* = 0.021), although not after Benjamini–Hochberg adjustment across 6 drivers (*Q* = 0.125). Because prevalence jointly reflects mutation supply and clonal selection, variation in one component may obscure or counteract variation in the other when prevalence is assessed along the colorectal continuum. We therefore examined these components separately.

**Table 2.** Is sidedness sufficient for prevalence, mutation supply, and cancer effect?

| Driver | Prevalence | Mutation supply | Cancer effect |
|---|---|---|---|
| *APC* | **0.021** | 0.734 | **0.019** |
| *PIK3CA* H1047 | 0.083 | **0.006** | 0.208 |
| *KRAS* codon 12/13 | 0.166 | 0.971 | 0.147 |
| *PIK3CA* E542/E545 | 0.285 | 0.327 | 0.339 |
| *BRAF* V600E | 0.714 | 0.305 | 0.875 |
| *TP53* | 0.863 | 0.227 | 0.772 |
| Total SNV burden | — | **0.010** | — |

Entries are *P* values comparing a side-only model against a colorectal-subsite-resolved model (Equations (2)–(7)); boldface indicates *P* < 0.05. All tests use *n* = 1027 whole-exome and -genome sequenced tumors except total SNV burden (whole-exome only, *n* = 1019). Tests carry 7 degrees of freedom.

*Mutation supply.* Tumor-specific neutral mutation rates *µ_i_* were analyzed on the logarithmic scale *y_i_*= log *µ_i_* using the Gaussian linear models

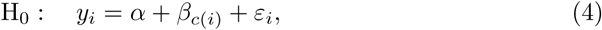

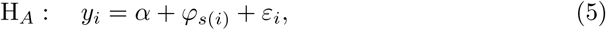

where *α* is the mean log mutation rate among all tumors; *β_c_*_(*i*)_ and *φ_s_*_(*i*)_ are the offsets of tumor *i*’s side and subsites from that grand mean, and the *ε_i_* are tumor-independent residuals. These nested models were compared using a partial *F* test with seven degrees of freedom.

As a complementary, driver-independent measure of overall mutational accumulation, we applied this same comparison to log-transformed total single-nucleotide variant burden *y_i_* = log *b_i_*, where *b_i_* is the total single-nucleotide variant count in tumor *i*. This analysis included the *n* = 1019 whole-exome sequenced tumors; eight whole-genome-sequenced tumors were excluded to avoid comparison of mutation counts obtained across different genomic coverage.

The side-only model was not rejected for five of six driver-specific neutral mutation rates (Table 2). The subsite-resolved model fit significantly better for *PIK3CA* H1047 (*P* = 0.006; Benjamini–Hochberg adjustment *Q* = 0.036) and total single-nucleotide variant burden (*P* = 0.010). Thus, both the supply of mutations capable of producing *PIK3CA* H1047 and overall mutational accumulation retain positional structure not represented by sidedness.

*Cancer effect.* Cancer effect was analyzed using cancereffectsizeR[33, 34], which models driver mutation status conditional on the tumor-specific mutation rate *µ_i_*and scaled selection coefficient *γ_i_*

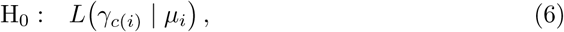

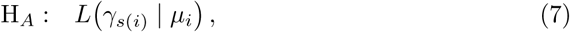

where *γ_i_* = 1 denotes neutrality, *γ_c_*_(*i*)_ is the maximum-likelihood estimate of selection on left-sided (*γ_L_*) and right-sided (*γ_R_*) tumors, and *γ_s_*_(*i*)_ is one value per subsite. The side-only and subsite-resolved models were

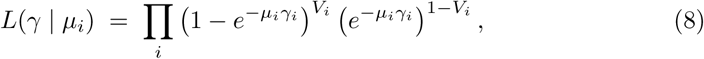

Neutral mutation rates were held fixed at their tumor-specific estimates, isolating spatial variation in selection. Maximized nested likelihoods were compared using a likelihood-ratio test, with seven degrees of freedom. The side-only model was not rejected for five of the six drivers (Table 2). *APC* showed nominal evidence of subsite variation in cancer effect (*P* = 0.019), although not after Benjamini–Hochberg adjustment (*Q* = 0.117).

Taken together, these analyses help explain why sidedness has endured: for many outcomes, the binary label captures much of the variation that current data can resolve. With current sample sizes, subsite resolution does not improve the fit of most driver-specific mutation-selection models, while the canonical features of right-sided disease—microsatellite instability, CIMP-high status, *BRAF* mutation, and serrated-pathway biology—form a coherent etiologic cluster. Even the cancer effect of *BRAF* V600E, which exhibited the strongest positional gradient among the drivers examined, was not significantly better explained by anatomical subsite than by side alone (*P* = 0.875). This result does not establish that each side is internally homogeneous, but it illustrates why sidedness remains clinically useful: it efficiently compresses several major biological signals into a binary variable.

Useful compression, however, is not biological equivalence. Dichotomization is hazardously expedient[56–58]: collapsing subsites into two sides can obscure nonmonotonic positional patterns. *APC*, the canonical gatekeeper of colorectal tumorigenesis, illustrates this potential failure. Its prevalence and cancer effect provide nominal evidence of subsite variation (*P* = 0.021, *P* = 0.019). *APC* prevalence increased across the left colorectum, declined at the splenic flexure, and increased again across the right. Collapsing these subsites into two sides averages away the opposing within-side changes, leaving little net left–right contrast. Yet treating nine subsites as unrelated categories costs seven degrees of freedom and sacrifices power. Continuous positional annotation, larger cohorts, and flexible spatial models should be investigated to distinguish genuine spatial structure from sampling variation.

The loss of positional information extends into the evidence used to guide treatment. Across pivotal trials in metastatic CRC, a binary side label serves as a primary analytic category: PARADIGM prespecified overall survival in the left-sided population as the first comparison in a hierarchical testing sequence, with the overall population tested only if it reached significance[14], while in CALGB/SWOG 80405 and in CRYSTAL and FIRE-3 the sidedness analyses were retrospective yet came to define how those trials are interpreted [7, 9]. In each, finer anatomical positioning is pooled into a single stratum. Clinical guidelines are affected by Clinical guidelines inherit this resolution: for *RAS* /*BRAF* wild-type, mismatch-repair-proficient disease, an anti-EGFR antibody is recommended for a left-sided primary and a bevacizumab-based regimen for a right-sided primary, with no treatment distinctions based on location within either side [1, 59], and little actual basis for decision in the case of transverse or splenic flexure tumors.

We need not continue to undertake Sysyphus’s unhappy labor. Unlike his blind prophetic neighbor in Hades, Tiresias, we cannot foretell what finer positional data will reveal. Yet current evidence shows that there are more sides to the colorectum than two. Rather than push a crude cutpoint back and forth, research cohorts, cancer registries, genomic repositories, and clinical trials should capture primary-tumor location at the finest practicable resolution. At minimum, this means retaining the ICD-O-3 subsite and, when feasible, distance from the nearest bowel landmark, recorded with the landmark, units, and measurement method. Harmonized collection would enable pooled analyses powered for clinically consequential inferences. Rectal cancer provides a precedent: distance from the anal verge is routinely recorded and guides management [27]. Extending fine-scale positional measurement throughout the colorectum would require structured capture during existing procedures, not a new assay or diagnostic intervention. The Sisyphean task of moving the cutpoint ends not when the perfect boundary is found, but when the boundary is no longer mistaken for anatomy.

## Supporting information

Supplementary Data and Codes

## List of Abbreviations

CALGB: Cancer and Leukemia Group B
CIMP: CpG island methylator phenotype
COSMIC: Catalogue of Somatic Mutations in Cancer
CRC: colorectal cancer
dMMR: deficient mismatch repair
EGFR: epidermal growth factor receptor
HPV: human papillomavirus
ICD-O-3: International Classification of Diseases for Oncology, Third Edition
MSI: microsatellite instability
OS: overall survival
SBS: single-base substitution
SNV: single-nucleotide variant
SWOG: Southwest Oncology Group

## Acknowledgment

The authors sincerely thank Yihan Liu and Kira A. Glasmacher of Department Computational Biology and Biomedical Informatics, Yale University for their contribution in initial brainstorming process of how to model subsites and the cancer effect.

## Declarations

### Ethics approval and consent to participate

Not applicable. This work is a perspective based exclusively on analyses of publicly available, de-identified datasets and does not involve new studies of human participants or collection of identifiable private information.

### Consent for publication

Not applicable. This manuscript does not contain any individual-level data, images, or other information requiring consent for publication.

### Data availability

The dataset analysed during the current study are available in the The Cancer Genome Atlas Research Network: https://www.cancer.gov/tcga and Catalogue of Somatic Mutations in Cancer: https://cancer.sanger.ac.uk/cosmic. Data filtering process can be found at **Supplementary Table S1, S2, and S3**, and **Supplementary Figure S1**.

### Materials availability

Not applicable.

### Code availability

Analytical tools can be found at https://townsend-lab-yale.github.io/cancereffectsizeR/. The code for this paper is available in supplementary information.

### Authors’ contributions

L.K. conceived the project, performed analyses, and composed the first draft. J.P.T. designed the analyses, supervised the work, and revised the manuscript. M.C. advised and revised the manuscript. And V.L.C. advised and revised the manuscript. All authors read and approved the final manuscript.

