## Supplementary Information for "Sisyphus at the splenic flexure: sidedness in colorectal cancer"

November 5, 2026

### 1 Supplementary methods

#### 1.1 Drivers selection

Each driver was analysed at the level at which its alterations constitute a distinct mechanistic entity, rather than uniformly at the gene level. For oncogenes activated by mechanistically distinct alleles, variants were separated accordingly; for tumor suppressors, deleterious variants were aggregated at the gene level, consistent with a loss-of-function mechanism in which any inactivating event is functionally equivalent. *BRAF* was analysed as the class-1 hotspot V600E (GENCODE v36 updates in GDC Data Release 32 denotes *BRAF* V640E[14]) only.Non-V600E *BRAF* drivers were classified following Yao et al. and Dankner et al.: class 2 (RAS-independent, dimer-forming: K601, L597, G464, and G469A/V) and class 3 (RAS-dependent, kinase-impaired: D594, G596, G466, and G469E). [1, 2]—occur in only about 2% of colorectal cancers [8], and in this cohort number 4 (class 2) and 9 (class 3), too infrequently to support subsite-resolved selection estimates. Other *BRAF* variants are unequivocally non drivers, which does not demand our considerations. *PIK3CA* was split into helical-domain (exon 9; E542K/E545K) and kinase-domain (exon 20; H1047R) mutations, which achieve gain of function by different mechanisms—helical mutants are *RAS*/*RTK*-dependent whereas kinase-domain mutants are *RAS*-independent [3, 4]. *KRAS* was grouped by hotspot codon into 12/13, 61 and 146, which differ in their biochemistry of *RAS* activation (GAP-insensitive P-loop mutants, mutants with impaired intrinsic and GAP-mediated hydrolysis, and fast-cycling nucleotide-exchange mutants, respectively) [5, 6]. However, codon 61 ($n = 13$) and 146 ($n = 30$) variants are too infrequent in many primary locations to support selection estimates. Therefore, we include in the supplementary methods just for bookkeeping. *NRAS* was analysed as a single hotspot group (Q61, G12, G13), parallel to *KRAS* but rarer ($n = 50$). Together with *KRAS* codons 61 and 146 and the non-V600E *BRAF* classes, *NRAS* was estimable only at the poles and is reported among the sparse driver groups (Table S7) rather than across all nine nine primary locations. *TP53* and *APC* were analysed as gene-level loss-of-function, aggregating all truncating and deleterious substitutions. The scaled selection coefficient (cancer effect) of each group was then estimated along the bowel axis with cancereffectsizeR [7]. *ERBB2* and *EGFR*, although part of *EGFR*-pathway signaling, act predominantly through amplification or overexpression rather than recurrent activating point mutations and are not assessable in a single-base-substitution cohort; they were therefore excluded. The final list of genes and variants of consideration is listed in Table S6.

#### 1.2 Driver prevalence analysis

For each driver group $g$ and subsite $s$, a tumor was scored a carrier if it harboured at least one qualifying variant, identified by genomic coordinate (hotspot windows for *BRAF* V600E, the *PIK3CA* helical and kinase domains, and *KRAS* codons 12/13, 61 and 146; gene-body loss-of-function for *APC* and *TP53*). The prevalence at subsite $s$ is $\hat{p}_{g,s} = k_{g,s}/n_s$, the number of carriers over the number of tumors with an assignable subsite ($N = 1{,}027$).

Because the mid-bowel nine primary locations are small (e.g. the splenic flexure, $n = 8$), uncertainty was quantified with the Wilson score 95% interval, which remains valid at small $n$ and near 0 or 1, rather than

| Subsite | ICD-O-3 | Length (cm) Mean | Length (cm) SD | Span (cm) Start–end | Distance to centre cm | Distance to centre Normalized |
|---|---|---|---|---|---|---|
| Rectum | C20 | 16.8 | 4.0 | 0.0 – 16.8 | 8.4 | 0.055 |
| Rectosigmoid[a] | C19 | — | | — | 16.8 | 0.110 |
| Sigmoid | C18.7 | 41.4 | 16.2 | 16.8 – 58.2 | 37.5 | 0.244 |
| Descending | C18.6 | 25.8 | 7.0 | 58.2 – 84.0 | 71.1 | 0.463 |
| Splenic flexure[a] | C18.5 | — | | — | 84.0 | 0.548 |
| Transverse | C18.4 | 46.6 | 9.6 | 84.0 – 130.6 | 107.3 | 0.699 |
| Hepatic flexure[a] | C18.3 | — | | — | 130.6 | 0.851 |
| Ascending | C18.2 | 22.8 | 4.4 | 130.6 – 153.4 | 142.0 | 0.926 |
| Cecum[a] | C18.0 | — | | — | 153.4 | 1.000 |

**Table S5.** Primary tumor location on a distance-from-anal-verge axis

Subsites are ordered by distance from the anal verge. Each ICD-O-3 subsite is placed at the distance from the anal verge to the centre of its segment, following Yamauchi et al.[9]. Segment lengths are the pooled adult means (mean ± SD) from Alqarni et al.[10], laid end to end from the anal verge (0 cm), so each span is measured from the anal verge; the axis length is the sum of the five means (153.4 cm; Alqarni's separately reported overall total is 148.3 cm).

the Wald interval:

$$\frac{\hat{p} + \dfrac{z^2}{2n} \;\pm\; z\sqrt{\dfrac{\hat{p}(1-\hat{p})}{n} + \dfrac{z^2}{4n^2}}}{1 + \dfrac{z^2}{n}}, \qquad z = 1.96.$$

Side-pooled prevalence (left, rectum–splenic flexure; right, transverse–cecum) were computed by summing carriers and tumors within each side and applying the same Wilson interval. Per-subsite prevalences (points, joined by a line, with Wilson whiskers) and side-pooled prevalences (horizontal bands) are displayed as in the established figure paradigm, with the clinically guided splenic-flexure cut-point marked.

### 1.3 Neutral mutation analysis

**Definition.** For each driver we quantified the *mutation supply*: the rate at which the driver's target site(s) would be mutated in a tumour in the *absence of selection*. This site-specific neutral (baseline) mutation rate is the denominator against which cancereffectsizeR [7] infers selection, and we use it here as a selection-free measure of supply. All rates were computed with cancereffectsizeR on the `ces.refset.hg38` reference.

**Computation.** The neutral rate of a single site in tumour $i$ factorises into a gene-level rate and a trinucleotide-context term,

$$\mu_i(\text{site}) \;=\; \underbrace{r_{\text{gene}}(i)}_{\text{gene rate}} \times \underbrace{\sum_k w_{ik}\, M_k\big(c(\text{site})\big)}_{\text{signature-weighted context rate}},$$

where $c$(site) is the site's trinucleotide context, $M_k$ is the COSMIC v3.4 definition of signature $k$ (probability of context $c$), and $w_{ik}$ are tumour $i$'s signature weights. The weights were obtained per tumour by mutational-signature deconvolution (`trinuc_mutation_rates`, signature set COSMIC v3.4, `sig_averaging_threshold` = 1, so that no tumour borrows the cohort-average composition); signatures were restricted to those expected in treatment-naive colorectal cancer (COSMIC exclusions for COAD/READ, retaining the POLE and mismatch-repair processes, i.e. not excluding SBS14). The gene-level rate $r_{\text{gene}}$ was estimated from tissue covariates (expression and replication timing; `gene_mutation_rates`). Site-level neutral rates were then read out with `baseline_mutation_rates`. For a driver *group* (a hotspot, codon set, or gene), the per-tumour supply is the sum of its constituent SNV sites, $\mu_i(\text{group}) = \sum_{\text{site}\in S} \mu_i(\text{site})$.

**Cohort.** Neutral rates require deconvolvable mutational spectra, so only whole-exome / whole-genome (WES/WGS) tumours were used; targeted panels were excluded. $\mu_i$ is defined for every WES/WGS tumour, whether or not it carries the driver.

**Table S6.** Driver groups analyzed and per-tumor carrier counts

| Group | $n$ | Driver mechanism |
|---|---|---|
| **Tumor suppressors** | | |
| *APC*[a] | 537 | Truncating loss-of-function; disrupts the $\beta$-catenin destruction complex $\rightarrow$ constitutive WNT signalling. |
| *TP53* | 530 | Loss-of-function ($\pm$ dominant-negative/gain-of-function hotspots); loss of the cell-cycle/apoptosis checkpoint, permitting genomic instability. |
| **Oncogenes** | | |
| *KRAS* codon 12/13 | 324 | P-loop mutants; block GAP-stimulated GTP hydrolysis; RAS locked GTP-bound $\rightarrow$ MAPK and PI3K. |
| *PIK3CA* helical (E542K/E545K, exon 9) | 91 | Gain-of-function by release of the p85 inhibitory contact; RAS/RTK-dependent; drives PI3K–AKT. |
| *BRAF* V600E | 64 | Class-1 constitutively active monomer; RAS-independent; hyperactivates MAPK (MEK–ERK). |
| *PIK3CA* kinase (H1047R, exon 20) | 42 | Gain-of-function by altered activation loop / membrane engagement; RAS-independent; drives PI3K–AKT. |

Carrier counts ($n$) are per-tumor, computed by genomic coordinate across the whole-exome/whole-genome analysis cohort ($n = 1{,}027$). Oncogenes are resolved to the variant or domain at which the activating mechanism differs; tumor suppressors are aggregated as gene-level loss-of-function. Within each category, groups are ordered by carrier count. [a]*APC* is undercounted: its canonical drivers are truncating frameshift indels, whereas this cohort is single-base-substitution focused, so only nonsense-substitution *APC* events are captured.

**Pooled estimation and the site set.** Crucially, $\mu$ was estimated from a *single* CESAnalysis over all WES/WGS tumours, rather than one analysis per subsite. The reason is the site set $S$: when a separate analysis is run per subsite, $S$ is restricted to the variants *observed* in that subsite, so a hyper-mutated (or larger) subsite observes more distinct sites and its summed $\mu$(group) is inflated—an artefact of sampling, not of mutability, that principally affects multi-site / whole-gene groups (*KRAS*, *APC*, *TP53*) and leaves single-hotspot groups (*BRAF* V600E) unaffected. Pooling fixes $S$ to the same cohort-wide set for every tumour, so the cross-subsite variation in $\mu$ reflects the mutational *process* (each tumour's own signatures) rather than the number of variants sampled. Per-tumour rates were averaged within subsites for display and analysis.

**Validation and limitations.** Because the mutational process is captured per tumor, subsite-specific supply is preserved by pooling; only tumours below the signature-averaging threshold are shifted from a subsite-average to a cohort-average composition. Two limitations are noted: (i) the reference provides only colon/rectum covariates, so gene-level mutability cannot vary continuously along the bowel and $\mu$'s finer spatial variation is carried entirely by the signature term; and (ii) for whole-gene groups the site set, though fixed, is the cohort-observed rather than the full theoretical target.

### 1.4 Attribution to signature analysis

Let variant $v$ have constituent SNVs $\mathcal{S}(v)$. For SNV $j$ (trinucleotide context $c_j$) carried by tumor $t$, the posterior probability that it was generated by signature $s$ is

$$\pi_{tjs} = \frac{w_{ts}\,\phi_s(c_j)}{\sum_{s'} w_{ts'}\,\phi_{s'}(c_j)}, \tag{1}$$

where $w_{ts}$ is tumor $t$'s weight on signature $s$ and $\phi_s(c)$ is signature $s$'s emission probability for context $c$ (mutational_signature_effects). The mutional signature $s$ is the mean tumor weight $\bar{w}_s = |T|^{-1} \sum_t w_{ts}$ (average_source_shares). Signatures were then collapsed into nine aetiological groups—defective mismatch repair, defective polymerase, defective homologous recombination, APOBEC, reactive oxygen species, base-excision-repair deficiency, two clock-like groups (SBS1; SBS5), and a residual non-actionable group—following Cannataro et al.[12], and renormalised to sum to one within each subsite. Figure 3 shows the mutation-supply and cancer-effect compositions across nine primary locations.

##### 1.4.1 Location-varying versus location-stable classification

Each aetiological group was labeled location-*varying* (drawn above the axis) or location-*stable* (below) by combining a significance test with an effect-size gate.

**Significance.** To avoid pseudo-replication, the test was performed at the *tumor* level. For tumors $t$ with SNV total $M_t$, the attributed count for group $g$ is $k_{tg} = \mathrm{round}\big(M_t \sum_{s \in g} w_{ts}\big)$. Negative-binomial generalized linear models with offset $\log M_t$ were fit and compared by likelihood-ratio test (LRT):

$$H_0: \ \log \mathbb{E}[k] = \beta_0 \qquad \text{versus} \qquad H_1: \ \log \mathbb{E}[k] = \beta_0 + \beta_1 x, \tag{2}$$

where $x$ is the normalized anal-verge coordinate, so $H_1$ is a monotone log-linear trend along the continuum (df = 1). The statistic $2(\ell_1 - \ell_0)$ was referred to $\chi^2_1$; the negative-binomial dispersion absorbs between-tumor heterogeneity.

### 1.5 Cancer effect size and its signature decomposition

#### 1.5.1 Per-subsite effect model

The cancer effect size $\gamma$ of a variant is a scaled selection coefficient: the factor by which selection multiplies the variant's flux to detectable frequency, relative to its neutral mutational baseline, so that $\gamma = 1$ is neutral and larger $\gamma$ means stronger positive selection [11]. For each sequenced tumor $j$ that is covered at the site, the baseline (neutral) rate at which the variant would arise is

$$\mu_j \ = \ r_{g(j)}\, q_j(c), \tag{3}$$

the product of the gene-level neutral mutation rate $r_{g(j)}$ (estimated by `dNdScv` with the colon/rectum covariates) and the tumor's relative rate $q_j(c)$ for the variant's specific trinucleotide substitution $c$ (from its deconvolved SBS weights). Selection scales this flux by $\gamma$, so under the model each tumor is an independent Bernoulli trial in which the variant is present with probability $1 - e^{-\gamma\mu_j}$ and absent with probability $e^{-\gamma\mu_j}$. Writing $\mathcal{P}$ for the covered tumors that carry the variant and $\mathcal{A}$ for those that do not, the effect size is the maximiser of the log-likelihood

$$\ell(\gamma) \ = \ \sum_{j \in \mathcal{P}} \log\big(1 - e^{-\gamma\mu_j}\big) \ - \ \sum_{j \in \mathcal{A}} \gamma\, \mu_j, \tag{4}$$

computed by `ces_variant`; the 95% confidence interval is the one-parameter likelihood-ratio (profile) interval on $\gamma$. A compound variant is scored the same way, with a tumor counted in $\mathcal{P}$ if it carries any constituent variant and $\mu_j$ summed over the constituents. Six driver groups matched the frequency panels: *BRAF* V600E; *KRAS* codon 12/13; *PIK3CA* helical (E542/E545/Q546); *PIK3CA* kinase (H1047/M1043/G1049); *APC*; and *TP53*. For *APC* and *TP53* every non-synonymous coding change (missense, nonsense, nonstop) and essential-splice variant was included, with no recurrence threshold (singletons retained) and synonymous changes excluded. Groups comprising more than one variant were merged into a single compound variant (define_compound_variants, unlimited merge distance). For each group the scaled selection intensity (cancer effect size) $\gamma$—the site-level maximum-likelihood ratio of the realised to the neutral mutational flux [11]—were obtained per subsite from the pooled fit described above. Because the design uses one indicator per subsite, the likelihood separates and each subsite's $\gamma$ is fitted on its own tumors alone; the 95% interval is the one-parameter profile-likelihood interval, $2(\hat{\ell} - \ell(\gamma)) = \chi^2_{1,0.95}$. All tumors in the subsite are used, whether or not they carry the driver.

#### 1.5.2 Signature decomposition of the effect

Each bar is filled with the SBS-source decomposition of that driver's effect, using the same method of Section 1.4. Because mutational_signature_effects accepts only individual (snv/aac) variant identifiers and not compound identifiers, the decomposition was computed over a group's *constituent* variants rather than its compound. Deconvolution again requires a deconvolvable spectrum, so it used only non-targeted (WES/WGS) tumors with $\geq 50$ SNVs; consequently a subsite shows no fill whenever none of its deconvolvable tumors carries the driver. Shares were collapsed into the same nine aetiological groups and renormalised per bar.

[13] J. G. Tate, S. Bamford, H. C. Jubb, Z. Sondka, D. M. Beare, N. Bindal, H. Boutselakis, C. G. Cole, C. Creatore, E. Dawson, *et al.*

[14] National Cancer Institute (NCI) Genomic Data Commons (GDC). GDC Data Release 32.0. Updated reference gene model to GENCODE v36 and re-harmonized genomic and transcriptomic pipelines. Available online at: https://gdc.cancer.gov/

COSMIC: the Catalogue of Somatic Mutations in Cancer. *Nucleic Acids Research*, 47(D1):D941–D947, 2019.

**Table S7.** Sparse driver groups along the anal-verge axis

| Subsite | $n$ | Carriers | Prevalence, % (95% CI) | Mutation supply | Cancer effect (95% CI) | Top 3 mutational sources |
|---|---|---|---|---|---|---|
| ***NRAS* Q61/G12/G13** (50 carriers) | | | | | | |
| Rectum | 382 | 17 | 4.5 (2.8 – 7.0) | $7.04 \times 10^{-6}$ | 1,727 (1,031 – 2,683) | clock SBS5 37%; unknown 29%; clock SBS1 11% |
| Rectosigmoid | 75 | 10 | 13.3 (7.4 – 22.8) | $3.26 \times 10^{-6}$ | 5,033 (2,521 – 8,832) | clock SBS5 42%; unknown 19%; dHR 18% |
| Sigmoid | 159 | 5 | 3.1 (1.4 – 7.1) | $2.44 \times 10^{-6}$ | 1,201 ( 431 – 2,582) | dHR 33%; clock SBS5 32%; dBER 15% |
| Descending | 19 | 2 | 10.5 (2.9 – 31.4) | $2.07 \times 10^{-6}$ | 4,223 ( 702 – 13,048) | dBER 36%; clock SBS5 25%; unknown 19% |
| Splenic flexure | 8 | 0 | 0.0 (0.0 – 32.4) | No observation | No observation | No observation |
| Transverse | 32 | 1 | 3.1 (0.6 – 15.7) | $8.52 \times 10^{-7}$ | 1,152 ( 66 – 5,073) | clock SBS5 60%; clock SBS1 17%; unknown 14% |
| Hepatic flexure | 14 | 0 | 0.0 (0.0 – 21.5) | No observation | No observation | No observation |
| Ascending | 130 | 1 | 0.8 (0.1 – 4.2) | $7.35 \times 10^{-6}$ | 275 ( 16 – 1,209) | dMMR 100%; clock SBS1 0%; dPOLE 0% |
| Cecum | 208 | 14 | 6.7 (4.1 – 11.0) | $2.41 \times 10^{-5}$ | 2,528 (1,424 – 4,094) | clock SBS5 38%; unknown 32%; clock SBS1 8% |
| ***KRAS* codon 146** (30 carriers) | | | | | | |
| Rectum | 382 | 9 | 2.4 (1.2 – 4.4) | $3.06 \times 10^{-6}$ | 2,554 (1,167 – 4,755) | dHR 31%; dBER 22%; clock SBS5 17% |
| Rectosigmoid | 75 | 2 | 2.7 (0.7 – 9.2) | $3.71 \times 10^{-7}$ | 4,166 ( 693 – 12,864) | clock SBS5 95%; unknown 5%; clock SBS1 0% |
| Sigmoid | 159 | 4 | 2.5 (1.0 – 6.3) | $2.53 \times 10^{-6}$ | 2,999 ( 931 – 6,969) | clock SBS5 40%; dMMR 25%; dBER 13% |
| Descending | 19 | 0 | 0.0 (0.0 – 16.8) | No observation | No observation | No observation |
| Splenic flexure | 8 | 0 | 0.0 (0.0 – 32.4) | No observation | No observation | No observation |
| Transverse | 32 | 2 | 6.2 (1.7 – 20.1) | $1.02 \times 10^{-5}$ | 3,620 ( 602 – 11,190) | dMMR 50%; clock SBS5 47%; unknown 3% |
| Hepatic flexure | 14 | 0 | 0.0 (0.0 – 21.5) | No observation | No observation | No observation |
| Ascending | 130 | 5 | 3.8 (1.7 – 8.7) | $7.20 \times 10^{-6}$ | 2,492 ( 894 – 5,357) | dMMR 45%; unknown 13%; dPOLE 12% |
| Cecum | 208 | 8 | 3.8 (2.0 – 7.4) | $2.84 \times 10^{-5}$ | 2,041 ( 877 – 3,948) | dMMR 39%; clock SBS5 24%; unknown 17% |
| ***KRAS* codon 61** (13 carriers) | | | | | | |
| Rectum | 382 | 6 | 1.6 (0.7 – 3.4) | $1.34 \times 10^{-6}$ | 2,047 ( 814 – 4,149) | unknown 63%; clock SBS5 12%; ROS 11% |
| Rectosigmoid | 75 | 0 | 0.0 (0.0 – 4.9) | No observation | No observation | No observation |
| Sigmoid | 159 | 0 | 0.0 (0.0 – 2.4) | No observation | No observation | No observation |
| Descending | 19 | 0 | 0.0 (0.0 – 16.8) | No observation | No observation | No observation |
| Splenic flexure | 8 | 0 | 0.0 (0.0 – 32.4) | No observation | No observation | No observation |
| Transverse | 32 | 0 | 0.0 (0.0 – 10.7) | No observation | No observation | No observation |
| Hepatic flexure | 14 | 0 | 0.0 (0.0 – 21.5) | No observation | No observation | No observation |
| Ascending | 130 | 2 | 1.5 (0.4 – 5.4) | $4.10 \times 10^{-6}$ | 2,026 ( 337 – 6,257) | dPOLE 48%; clock SBS5 22%; unknown 21% |
| Cecum | 208 | 5 | 2.4 (1.0 – 5.5) | $8.71 \times 10^{-6}$ | 3,659 (1,312 – 7,864) | clock SBS5 48%; unknown 22%; dMMR 18% |
| ***BRAF* non-V600E class 3** (9 carriers) | | | | | | |
| Rectum | 382 | 2 | 0.5 (0.1 – 1.9) | $1.20 \times 10^{-6}$ | 750 ( 125 – 2,316) | clock SBS5 71%; clock SBS1 17%; unknown 12% |
| Rectosigmoid | 75 | 0 | 0.0 (0.0 – 4.9) | No observation | No observation | No observation |
| Sigmoid | 159 | 3 | 1.9 (0.6 – 5.4) | $2.60 \times 10^{-6}$ | 2,512 ( 625 – 6,515) | unknown 37%; clock SBS5 27%; dBER 17% |
| Descending | 19 | 0 | 0.0 (0.0 – 16.8) | No observation | No observation | No observation |
| Splenic flexure | 8 | 0 | 0.0 (0.0 – 32.4) | No observation | No observation | No observation |
| Transverse | 32 | 0 | 0.0 (0.0 – 10.7) | No observation | No observation | No observation |
| Hepatic flexure | 14 | 0 | 0.0 (0.0 – 21.5) | No observation | No observation | No observation |
| Ascending | 130 | 1 | 0.8 (0.1 – 4.2) | $1.57 \times 10^{-6}$ | 1,252 ( 71 – 5,513) | dPOLE 79%; dMMR 21%; clock SBS1 0% |
| Cecum | 208 | 3 | 1.4 (0.5 – 4.2) | $6.71 \times 10^{-6}$ | 1,587 ( 264 – 4,901) | unknown 46%; APOBEC 43%; clock SBS5 6% |

*No observation* marks a subsite where the quantity could not be estimated: cancer effect and mutation supply require at least one carrier there, and the mutational sources additionally require carriers among tumors with ≥50 SNVs. Driver groups are ordered by total carriers (given beside each group name); subsites run from rectum to cecum. All analyses use the WES/WGS data ($n = 1027$ tumors).

| Subsite | $n$ | Carriers | Prevalence, % (95% CI) | Mutation supply | Cancer effect (95% CI) | Top 3 mutational sources |
|---|---|---|---|---|---|---|
| ***BRAF* non-V600E class 2** (4 carriers) | | | | | | |
| Rectum | 382 | 0 | 0.0 (0.0 – 1.0) | No observation | No observation | No observation |
| Rectosigmoid | 75 | 0 | 0.0 (0.0 – 4.9) | No observation | No observation | No observation |
| Sigmoid | 159 | 1 | 0.6 (0.1 – 3.5) | $1.21 \times 10^{-6}$ | 958 ( 55 – 4,216) | unknown 89%; ROS 11%; clock SBS1 0% |
| Descending | 19 | 0 | 0.0 (0.0 – 16.8) | No observation | No observation | No observation |
| Splenic flexure | 8 | 0 | 0.0 (0.0 – 32.4) | No observation | No observation | No observation |
| Transverse | 32 | 0 | 0.0 (0.0 – 10.7) | No observation | No observation | No observation |
| Hepatic flexure | 14 | 0 | 0.0 (0.0 – 21.5) | No observation | No observation | No observation |
| Ascending | 130 | 1 | 0.8 (0.1 – 4.2) | $1.83 \times 10^{-6}$ | 1,515 ( 86 – 6,669) | No observation |
| Cecum | 208 | 2 | 1.0 (0.3 – 3.4) | $7.75 \times 10^{-6}$ | 2,897 (720 – 7,512) | clock SBS5 74%; ROS 15%; unknown 6% |

**Table S7** (continued). Sparse driver groups along the anal-verge axis

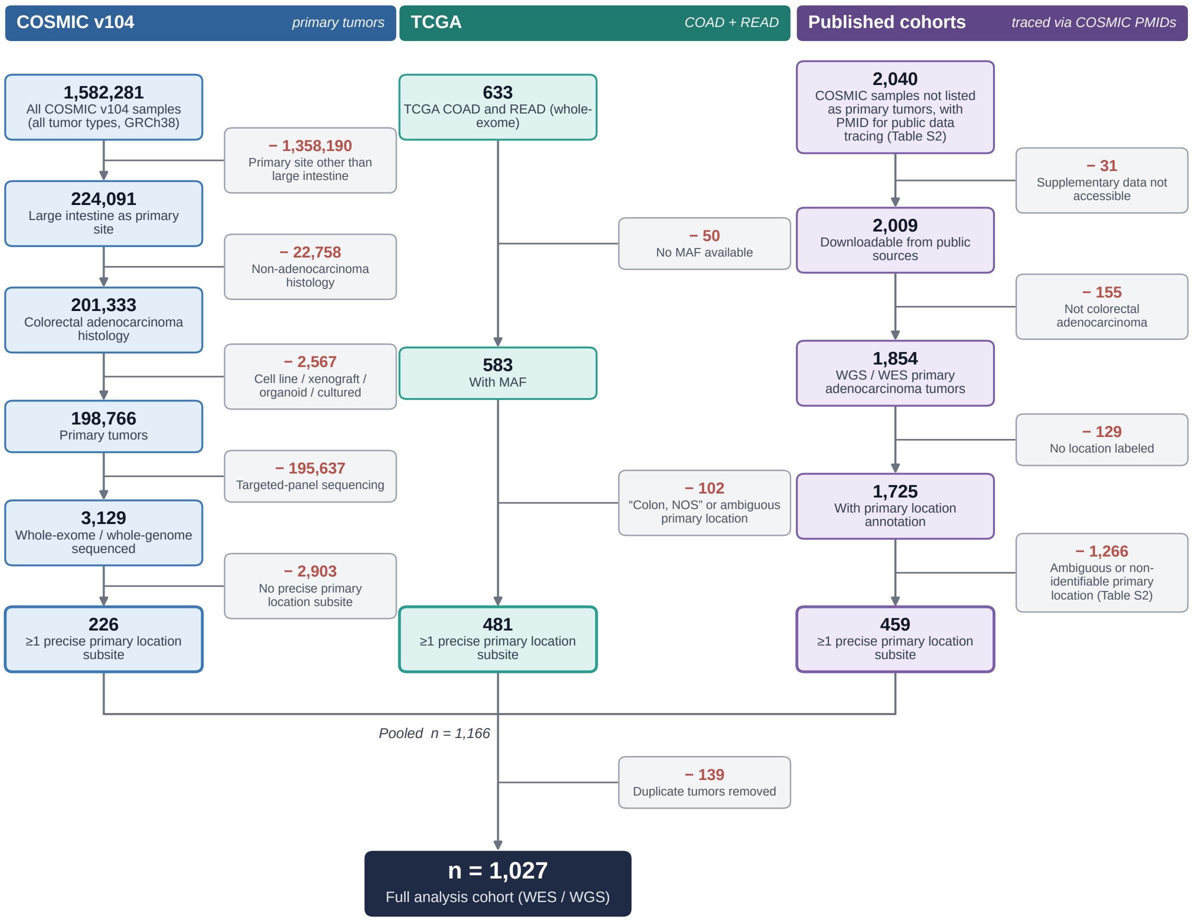


**Figure S1. Assembly pipeline of the analysis cohort.** Pipeline of sample selection from COSMIC v104[13] and TCGA COAD/READ to the $n$ = 1,027-cohort; filled boxes are retained counts, shaded boxes the number removed and the reason. The COSMIC discovery yield 226 whole-exome/whole-genome. TCGA contributes 481 subsite-precise cases, and publication recovery of PMID-traceable COSMIC samples (see **Supplementary Tables S1-S4**) contributes 459. Merging the four streams and removing 139 duplicates yields the final cohort. All analyses in this work use the whole-exome/whole-genome-sequenced tumors ($n$ = 1,027).
